# Next-generation Sequence-analysis Toolkit (NeST): A standardized bioinformatics framework for analyzing Single Nucleotide Polymorphisms in next-generation sequencing data

**DOI:** 10.1101/323535

**Authors:** Shashidhar Ravishankar, Sarah E. Schmedes, Dhruviben S. Patel, Mateusz Plucinski, Venkatachalam Udhayakumar, Eldin Talundzic, Fredrik Vannberg

## Abstract

Rapid advancements in next-generation sequencing (NGS) technologies have led to the development of numerous bioinformatics tools and pipelines. As these tools vary in their output function and complexity and some are not well-standardized, it is harder to choose a suitable pipeline to identify variants in NGS data. Here, we present NeST (NGS-analysis Toolkit), a modular consensus-based variant calling framework. NeST uses a combination of variant callers to overcome potential biases of an individual method used alone. NeST consists of four modules, that integrate open-source bioinformatics tools, a custom Variant Calling Format (VCF) parser and a summarization utility, that generate high-quality consensus variant calls. NeST was validated using targeted-amplicon deep sequencing data from 245 *Plasmodium falciparum* isolates to identify single-nucleotide polymorphisms conferring drug resistance. The results were verified using Sanger sequencing data for the same dataset in a supporting publication [28]. NeST offers a user-friendly pipeline for variant calling with standardized outputs and minimal computational demands for easy deployment for use with various organisms and applications.

## Introduction

Innovations in sequencing technology have led to a massive increase in throughput coupled with a rapid decrease in cost per base sequenced. As a result, speed and accuracy of next generation sequencing (NGS) have improved exponentially in the past few years [21, 22]. The vast increase in the number of NGS platforms and applications has led to the development of myriad bioinformatics tools to perform standard analyses, such as variant calling [23, 30]. Many bioinformatics pipelines have been developed to incorporate variant calling tools to ensure that only high quality variant calls are reported [3, 9, 12, 25]; however, these solutions are mainly tailored for model organisms [5, 7]. Modified approaches are required for organisms that lack extensive curated databases and complete annotations. Here, we introduce NeST (Next-generation Sequence-analysis Toolkit), a consensus-based variant calling framework that integrates multiple variant calling strategies to improve the accuracy of variant call. The plug-and-play framework integrates standard variant callers such as GATK [8] and Samtools [17] and allows users to plug-in their tools of choice to the existing consensus system. To standardize and simplify the analysis process a standard BED file format is used for annotation. The tab delimited BED (Browser Extensible Format) file format is easy to generate and annotations in BED format can be easily accessed, including via UCSC table browser [13]. NeST was validated using targeted-amplicon deep sequencing (TADS) data from *Plasmodium falciparum*, to accurately identify variants that are associated with resistance to anti-malarial treatments, as compared with Sanger sequencing data [28]. The results were verified using Sanger sequencing technology in a previously published study [28]. The framework has been tested on Linux and OSX operating systems for deployment across these two platforms.

## Analysis framework

NeST implements a consensus-based variant calling framework that integrates open-source bioinformatics tools for quality correction, alignment, and variant calling from NGS datasets. The ease of deployment and accuracy is improved by incorporation of four custom modules that simplify the process of variant calling. The modules are described below and illustrated in Fig 1.

**Figure 1.**
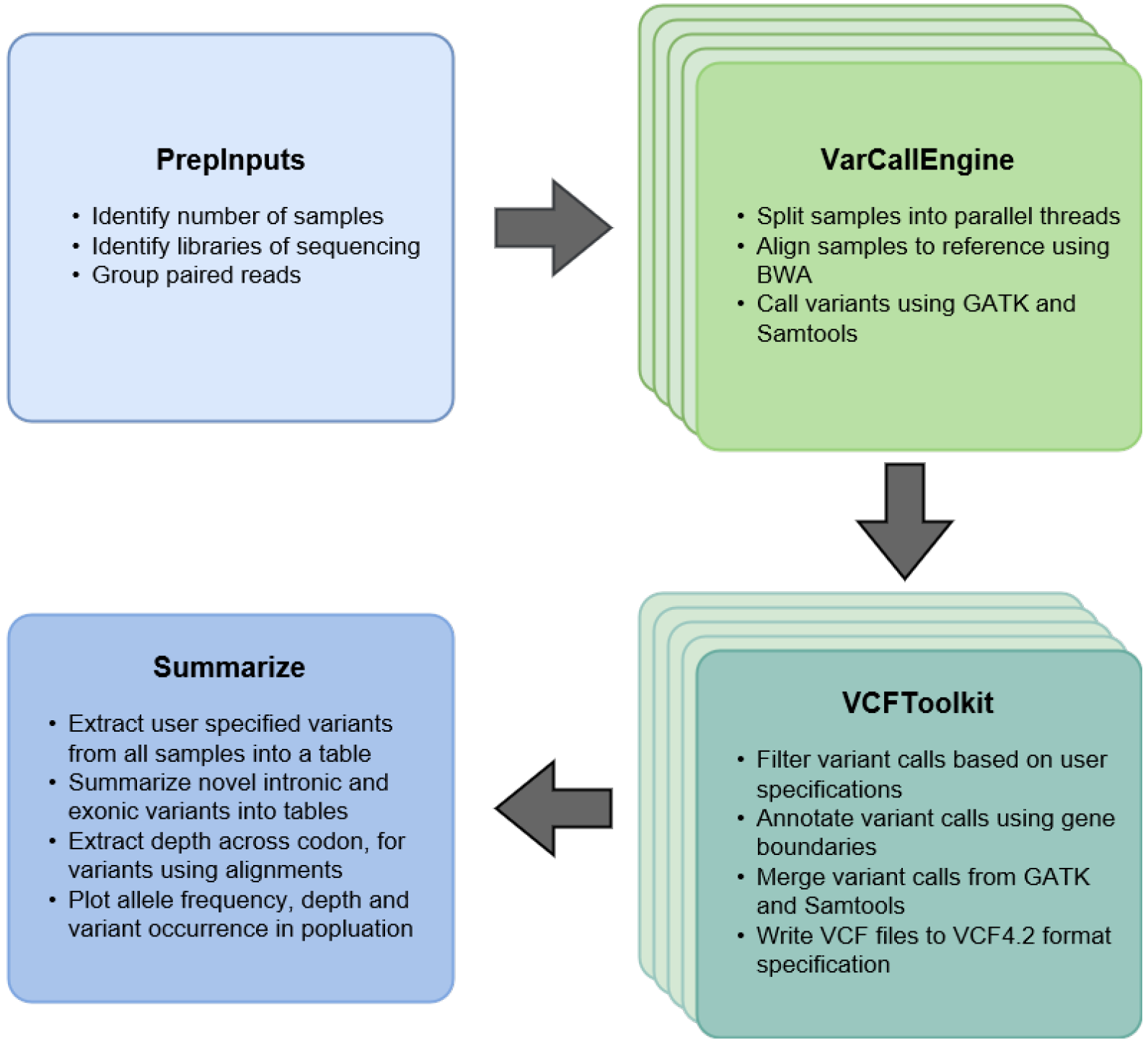
The four key modules of NeST. Multiple instances of VarCallEngine and VCFToolkit, are spawned into parallel processes to analyze each sample in the dataset

### PrepInputs Module

The module takes one of two formats as input, either an SRA (Sequence Read Archive [15]) accession list or a path to a directory containing sequence data in Fastq format. If the input is an SRA accession list, Fastq files are retrieved from the SRA using SRAToolkit [15]. Alternatively, if the input is a path to a directory, the PrepInputs module scans through each level of the directory to retrieve all Fastq files present within the parent directory and each subdirectory. The module parses each Fastq file and uses the Fastq headers to retrieve the relevant sequencing run information, including sample name, library type, and sequence length. Fastq files are grouped by sample name, and then a run dictionary is created for the study. The samples are then broken into parallel processes for further analysis. For default settings, four parallel threads are spawned to analyze four samples at one time.

### VarCallEngine Module

Two sets of variants calls are generated for each sample by the VarCallEngine. First, sequence reads are trimmed and cleaned according to quality thresholds, and adapters are removed using BBDuk [4]. To reduce sequence artifacts and ensure high-quality alignments, reads with a quality score less than 30 are trimmed and reads less than 100 bases in length are removed prior to analysis. Cleaned sequence reads are then aligned to a reference genome using one of the four available aligners (BWA, Bowtie2, BBMap, and SNAP [4, 16, 18, 31]), which can be chosen by the user. If none are specified, BWA is used to align the reads under default settings. All aligners are run in global alignment mode. The alignments are then sorted, de-duplicated and read group information is added using Samtools [19] and Picard tools [1]. The de-duplicated BAM files are then used to call variants using GATK [8] and Samtools-Bcftools [17]. The current framework restricts calls to SNPs and ignores short Inserts and Deletions(InDels).

### VCFToolkit Module

The VCFToolkit is a custom VCF parser to filter, merge, and annotate variant calls from GATK [8] and Samtools [17]. The parser is broken into submodules, each built to perform a function, allowing for ease of use and addition of extra functionality as per user requirements. The Filter module allows the user to filter variants by three categories, QUAL (quality) filter, INFO (information) filter and FORMAT filter. The Filter module follows the VCF conventions regarding the input of QUAL, INFO and FORMAT filters. The filters are accepted as a string of key-value pairs for the INFO and FORMAT field, and a float for the QUAL field, as denoted in a VCF file and delimited by a semi-colon when specifying more than one key. The filtered variant calls are then merged using the Merge module. This module sequentially parses through the VCF files and merges all headers, INFO and FORMAT field values. Additionally, the module adds an INFO field annotation called “Conf” (short for Confidence), indicating the number of variant callers that the SNP was found in. The VCF files from the variant callers and Merge module are then annotated using a BED file, with designated gene boundaries, provided by the user. The gene boundaries file is in 12 column BED format, with one gene per line of the file and exon coordinates specified for each gene. This light-weight annotation tool allows for easier annotation of VCF files from organisms that lack a comprehensive annotated database.

### Summarize Module

To enable the comparison of results across all samples in the study, the annotated variants from VCF files for each sample are extracted and grouped into summary tables using the Summarize module. A data-frame is created summarizing the occurrence of user provided reportable variants within all samples. To track novel variants that were not included in the reportable variant list, novel exonic and intronic variants are grouped into two separate tables (Supplementary File S2, S3, S4). For each variant location, the alignment files are scanned to extract the depth at a codon level to help distinguish between a variant and wild-type calls. Visualizations depicting allele frequency, sequencing depth, and occurrence of variants across all samples are generated using custom RScripts provided with NeST (Supplementary File S1).

The source and installation instructions for NeST can be found on Github (https://github.com/shashidhar22/NeST.git).

## Framework Evaluation

NeST was developed to enable rapid and accurate drug resistance analysis of *P. falciparum*. The current version of NeST has been implemented in the Malaria Resistance Surveillance (MaRS) pipeline developed at the Centers for Disease Control and Prevention(CDC) [28] and the modular nature of the framework can easily be adapted for variant identification with other pathogens. The performance of NeST was tested on an MSI GLM62M laptop, 8GB RAM and Intel i5 processor, on Ubuntu 16.04. Targeted-Amplicon Deep Sequencing (TADS) data (i.e., *crt*, *mdrl*, *k13*, *dhfr*, *dhps*, and *cytochrome b* genes and mitochondrial genome) from 245 *P. falciparum* samples, generated from MaRS [28], was analyzed in 1 hour and 3 minutes. All variant calls were independently validated using the Geneious [14] software package. Of the 26 drug resistant SNPs in this MaRS dataset, Geneious and NeST both identified 1935 instances of these SNPs and 4237 instances were wild type alleles, in the 245 samples (BioProject PRJNA428490). NeST identified an additional 169 instances of these SNPs not found by Geneious. In comparison, Geneious identified 29 additional instances of these SNPs missed by NeST (Supplementary File S1, S5, S6).

## Conclusion

Although variant calling has been a standard analysis for NGS data, numerous studies have shown that many tools do not always produce concordant results [2, 6, 10, 20, 23, 24, 26, 29, 32]. Inaccurate or discordant variant calling can greatly affect interpretation of results and have wide implications for the validity of clinical and/or public health-oriented program related data including surveillance data and diagnosis. Efforts are being made to standardize and validate pipelines to improve the accuracy and reproducibility of results; however, it is becoming evident that specific bioinformatics analysis pipelines may need to be developed and/or evaluated for specific organisms [3, 5, 7, 9, 11, 27]. Here, we present a variant calling python framework, NeST, that can be used to identify known and novel polymorphisms and ensure reproducibility of annotated variant calls.

## References

1. Picard tools https://broadinstitute.github.io/picard, 2018.

2. D. C. Bauer. Variant calling comparison. Brain, 1, 2011.

3. R. W. W. Brouwer, M. C. G. N. Van den hout, F. G. Grosveld, and W. F. J. Van ijcken. NARWHAL, a primary analysis pipeline for NGS data. Bioinformatics, 2012.

4. B. Bushnell. BBMap: a fast, accurate, splice-aware aligner. Technical report, 2014.

5. M. Chiara, S. Gioiosa, G. Chillemi, M. D’Antonio, T. Flati, E. Picardi, F. Zambelli, D. S. Horner, G. Pesole, and T. Castrignanò. CoVaCS: a consensus variant calling system. BMC Genomics, 19(1):120, 2018.

6. A. Cornish and C. Guda. A Comparison of Variant Calling Pipelines Using Genome in a Bottle as a Reference. BioMed Research International, 2015, 2015.

7. M. D’Antonio, P. D’Onorio De Meo, D. Paoletti, B. Elmi, M. Pallocca, N. Sanna, E. Picardi, G. Pesole, and T. Castrignanò. WEP: a high-performance analysis pipeline for whole-exome data. BMC Bioinformatics, 2013.

8. M. A. DePristo, E. Banks, R. Poplin, K. V. Garimella, J. R. Maguire, C. Hartl, A. A. Philippakis, G. del Angel, M. A. Rivas, M. Hanna, A. McKenna, T. J. Fennell, A. M. Kernytsky, A. Y. Sivachenko, K. Cibulskis, S. B. Gabriel, D. Altshuler, and M. J. Daly. A framework for variation discovery and genotyping using next-generation DNA sequencing data. Nature Genetics, 43(5):491–498, 2011.

9. K. M. Fisch, T. Meißner, L. Gioia, J. C. Ducom, T. M. Carland, S. Loguercio, and A. I. Su. Omics Pipe: A community-based framework for reproducible multi-omics data analysis. Bioinformatics, 31(11):1724–1728, 2015.

10. E. Giannoulatou, S.-H. Park, D. T. Humphreys, and J. W. Ho. Verification and validation of bioinformatics software without a gold standard: a case study of BWA and Bowtie. From Asia Pacific Bioinformatics Network (APBioNet) Thirteenth International Conference on Bioinformatics, 2014.

11. J. Goecks, A. Nekrutenko, J. Taylor, and T. Galaxy Team. Galaxy: a comprehensive approach for supporting accessible, reproducible, and transparent computational research in the life sciences. Genome Biology, 2010.

12. B. N. Howie, P. Donnelly, and J. Marchini. A flexible and accurate genotype imputation method for the next generation of genome-wide association studies. PLoS Genetics, 5(6), 2009.

13. D. Karolchik. The UCSC Table Browser data retrieval tool. Nucleic Acids Research, 32(90001):493D–496, 2004.

14. M. Kearse, R. Moir, A. Wilson, S. Stones-Havas, M. Cheung, S. Sturrock, S. Buxton, A. Cooper, S. Markowitz, C. Duran, T. Thierer, B. Ashton, P. Meintjes, and A. Drummond. Geneious Basic: An integrated and extendable desktop software platform for the organization and analysis of sequence data. Bioinformatics, 2012.

15. Y. Kodama, M. Shumway, and R. Leinonen. The sequence read archive: Explosive growth of sequencing data. Nucleic Acids Research, 40(D1):2011–2013, 2012.

16. B. Langmead and S. L. Salzberg. Fast gapped-read alignment with Bowtie 2. Nature methods, 9(4):357–9, 2012.

17. H. Li. A statistical framework for SNP calling, mutation discovery, association mapping and population genetical parameter estimation from sequencing data. Bioinformatics, 27(21):2987–2993, 2011.

18. H. Li. Aligning sequence reads, clone sequences and assembly contigs with BWA-MEM. 00(00):1–3, 2013.

19. H. Li and R. Durbin. Fast and accurate short read alignment with Burrows ‑ Wheeler transform. Bioinformatics, 25(14):1754–1760, 2009.

20. X. Liu, S. Han, Z. Wang, J. Gelernter, and B. Z. Yang. Variant Callers for Next-Generation Sequencing Data: A Comparison Study. PLoS ONE, 8(9), 2013.

21. E. R. Mardis. A decade’s perspective on DNA sequencing technology. Nature, 470(7333):198–203, 2011.

22. E. R. Mardis. DNA sequencing technologies: 2006-2016. Nature Protocols, 12(2):213–218, 2017.

23. J. O’Rawe, T. Jiang, G. Sun, Y. Wu, W. Wang, J. Hu, P. Bodily, L. Tian, H. Hakonarson, W. E. Johnson, Z. Wei, K. Wang, and G. J. Lyon. Low concordance of multiple variant-calling pipelines: practical implications for exome and genome sequencing. Genome Medicine, 2013.

24. S. Pabinger, A. Dander, M. Fischer, R. Snajder, M. Sperk, M. Efremova, B. Krabichler, M. R. Speicher, J. Zschocke, and Z. Trajanoski. A survey of tools for variant analysis of next-generation genome sequencing data. Briefings in Bioinformatics, 15(2):256–278, 2014.

25. J. Reumers, P. De Rijk, H. Zhao, A. Liekens, D. Smeets, J. Cleary, P. Van Loo, M. Van Den Bossche, K. Catthoor, B. Sabbe, E. Despierre, I. Vergote, B. Hilbush, D. Lambrechts, and J. Del-Favero. Optimized filtering reduces the error rate in detecting genomic variants by short-read sequencing. Nature Biotechnology, 30(1), 2011.

26. N. Rieber, M. Zapatka, B. Rbel Lasitschka, D. Jones, P. Northcott, B. Hutter, N. Jä Ger, M. Kool, M. Taylor, P. Lichter, S. Pfister, S. Wolf, B. Brors, R. Eils, and O. Hofmann. Coverage Bias and Sensitivity of Variant Calling for Four Whole-genome Sequencing Technologies. 2013.

27. J. Singer, H. J. Ruscheweyh, A. L. Hofmann, T. Thurnherr, F. Singer, N. C. Toussaint, C. K. Ng, S. Piscuoglio, C. Beisel, G. Christofori, R. Dummer, M. N. Hall, W. Krek, M. P. Levesque, M. G. Manz, H. Moch, A. Papassotiropoulos, D. J. Stekhoven, P. Wild, T. Wüst, B. Rinn, and N. Beerenwinkel. NGS-pipe: A flexible, easily extendable and highly configurable framework for NGS analysis. Bioinformatics, 34(1):107–108, 2018.

28. E. Talundzic, S. Ravishankar, J. Kelly, D. Patel, M. Plucinski, S. Schmedes, D. Ljolje, B. Clemons, S. Madison-Antenucci, P. M. Arguin, N. Lucchi, F. Vannberg, and V. Udhayakumar. A next-generation sequencing and bioinformatics protocol for Malaria drug Resistance marker Surveillance (MaRS). Antimicrobial Agents and Chemotherapy, (February):AAC.02474–17, 2018.

29. A. Talwalkar, J. Liptrap, J. Newcomb, C. Hartl, J. Terhorst, K. Curtis, M. Bresler, Y. S. Song, M. I. Jordan, and D. Patterson. SMaSH: A benchmarking toolkit for human genome variant calling. Bioinformatics, 30(19):2787–2795, 2014.

30. H. Xu, J. DiCarlo, R. Satya, Q. Peng, and Y. Wang. Comparison of somatic mutation calling methods in amplicon and whole exome sequence data. BMC Genomics, 15(1): 244, 2014.

31. M. Zaharia, W. J. Bolosky, K. Curtis, A. Fox, D. Patterson, S. Shenker, I. Stoica, R. M. Karp, and T. Sittler. Faster and More Accurate Sequence Alignment with SNAP.

32. J. M. Zook, B. Chapman, J. Wang, D. Mittelman, O. Hofmann, W. Hide, and M. Salit. Integrating human sequence data sets provides a resource of benchmarking SNP and indel genotype calls. Nature Biotechnology, 32(3):246–251, 2014.

